# A rapid F0 CRISPR screen in zebrafish to identify regulators of neuronal development in the enteric nervous system

**DOI:** 10.1101/2021.07.17.452230

**Authors:** Ann E Davidson, Nora RW Straquadine, Sara A Cook, Christina G Liu, Julia Ganz

## Abstract

The enteric nervous system (ENS) provides the intrinsic innervation of the gastrointestinal (GI) tract with millions of neurons and diverse neuronal subtypes and glial cells. The ENS regulates essential gut functions such as motility, nutrient uptake, and immune response, but basic information about the genes that control ENS neuronal specification and differentiation remains largely unknown. Deficits in ENS neuron numbers and composition cause gut dysfunction with debilitating GI symptoms, and are associated with e.g. Hirschsprung disease, inflammatory gut diseases, autism spectrum disorder, and neurodegenerative diseases such as Parkinson’s disease. The genetic basis of most of these ENS disorders remains unknown. Recent transcriptomic analyses have identified many candidate genes for regulating ENS neurogenesis. However, functional evaluation of these candidate genes significantly lags because experimental testing of their role in ENS neurogenesis is time-consuming and expensive. Here, we have developed a rapid, scalable F_0_ CRISPR genome editing screen in zebrafish to functionally determine which candidate genes control neuronal development in the ENS. Proof-of-concept experiments targeting the known ENS regulators *sox10* and *ret* phenocopy stable mutants with high efficiency and precision showing that our approach is reliable to identify regulators ENS neurogenesis using F_0_ guide RNA-injected larvae (F_0_ crispants). We then evaluate the role of 10 transcription factor genes for regulating ENS neurogenesis and function. Pools of guide RNAs targeting 2-3 candidate genes are co-injected with Cas9 protein into one-cell stage *phox2bb:GFP* transgenic zebrafish embryos to directly assess qualitative change in ENS neuron numbers compared to controls in 6-day old F_0_ crispants. Target genes from crispant pools exhibiting reduced ENS neuronal numbers were then tested individually to identify the responsible gene(s). We identify five transcription factors that show a reduction in ENS neurons indicating an influence on enteric progenitor cell differentiation into ENS neurons. Adding a simple and efficient test to further assess crispant gut motility, we find that loss-of-function of two of the transcription factor genes reduced intestinal transit of fluorescently labeled food through the gut. In summary, our novel, multistep, yet straight-forward CRISPR screening approach in zebrafish enables testing the genetic basis of ENS developmental and disease gene functions that will facilitate high-throughput evaluation of the manifold candidate genes emerging from transcriptomic, genome-wide association or other ENS-omics studies. Such *in vivo* ENS crispant screens will contribute to a better understanding of ENS neuronal development regulation in vertebrates and what goes awry in ENS disorders.

## Introduction

A large proportion of the general population in Western countries suffer from unsolved gastrointestinal (GI) symptoms drastically affecting quality of life. Diseases such as Hirschsprung disease, autism spectrum disorder, and inflammatory gut diseases affect the bowel, present with debilitating GI symptoms, and are associated with enteric nervous system (ENS) dysfunction ^1, 2^. The ENS innervates the GI tract and controls all essential gut functions, such as motility and nutrient uptake ^3, 4^. Consequently, abnormal ENS neurogenesis impairs gut function. Because few genes that regulate ENS neurogenesis have thus far been identified ^3, 4^, the genetic basis of many ENS disorders and gut diseases with an ENS component are not known. A recent surge in transcriptomic data has generated a wealth of novel, but functionally untested candidate genetic regulators of ENS neurogenesis ^2, 4, 5^.

Here, we present a rapid and cost-effective CRISPR/Cas9-based reverse genetic screening pipeline to functionally test important candidate regulators of ENS neurogenesis in zebrafish. Zebrafish digestive tract development, anatomy, and physiology is very similar to mammals. Additionally, the small, externally developing zebrafish larvae exhibit a fully functional and transparent gut by 5 days post fertilization (dpf), allowing for the rapid evaluation of ENS phenotypes and GI function ^3, 4^.

## Results

First, F_0_ guide RNA-injected zebrafish larvae (crispants) were examined for qualitative reduction in enteric neuron number. As proof-of-principle, we co-injected Cas9 protein and guide RNAs (gRNAs) targeting the known ENS developmental regulators *sox10* or *ret* in *phox2bb:EGFP* transgenic zebrafish larvae (referred to as *phox2bb:GFP*^+^). At larval stages, *phox2bb:GFP*^+^ cells are mostly ENS neurons with a small number of enteric progenitor cells ^6^. Stable mutants of *sox10* and *ret* completely lack enteric neurons ^3, 4^. Likewise, *sox10* and *ret* F_0_ crispants show a complete absence of ENS neurons (Suppl Fig. 1A-C, F,G). Both *sox10* and *ret* crispants exhibit robust efficiency of Insertion and/or Deletions (InDels) at the target site (Suppl Fig. 1H). A gRNA targeting the *slc24a5* gene is coinjected to provide a positive visual control ^7^. At 2 days, *slc24a5* crispants have a highly consistent reduction of pigmentation (*golden* phenotype) allowing for quick visual screening of injection efficiency (Suppl Fig. 1D,E).

To identify new regulators of ENS neurogenesis, 10 target candidate genes (Suppl Table 1) were selected from previous transcriptome analyses in the ENS ^5^. These genes were selected because we aimed to test our CRISPR screening approach on genes predicted to regulate different aspects of ENS neurogenesis, e.g. stem cell maintenance, neuronal specification, and differentiation. For higher screening efficiency, we grouped these genes in four pools targeting 2-3 genes per pool (Suppl Table 1). Each pool consists of genes with a similar proposed role in ENS neurogenesis and we grouped duplicate genes (paralogs) in one pool, e.g. *jarid2a* and *jarid2b*. Following the steps of the CRISPR screen as outlined in Fig 1, we screened for changes in *phox2bb:GFP*^+^ cells at 6 days when ENS neurons are well-established. A proportion of crispants have fewer GFP^+^ cells along the gut in all pools except pool 2 (Figure 1A-H). For all genes, we found robust efficiency of 64-91% InDels in crispant genomic DNA and good survival after injection (Fig. 1I, Suppl Fig 1I). To test which of the gene(s) are responsible for the ENS phenotypes, each target gene gRNA was then injected individually, except for *homeza* and *homezb* because they did not show an ENS phenotype in the pool injection. We found that functional loss of *jarid2a*, *mycn*, *foxj3*, *dlx1a*, and *phox2a* results in fewer GFP^+^ cells, whereas targeting *foxj2, foxn3* and *jarid2b* does not consistently reduce GFP^+^ cell numbers.

**Fig 1.**
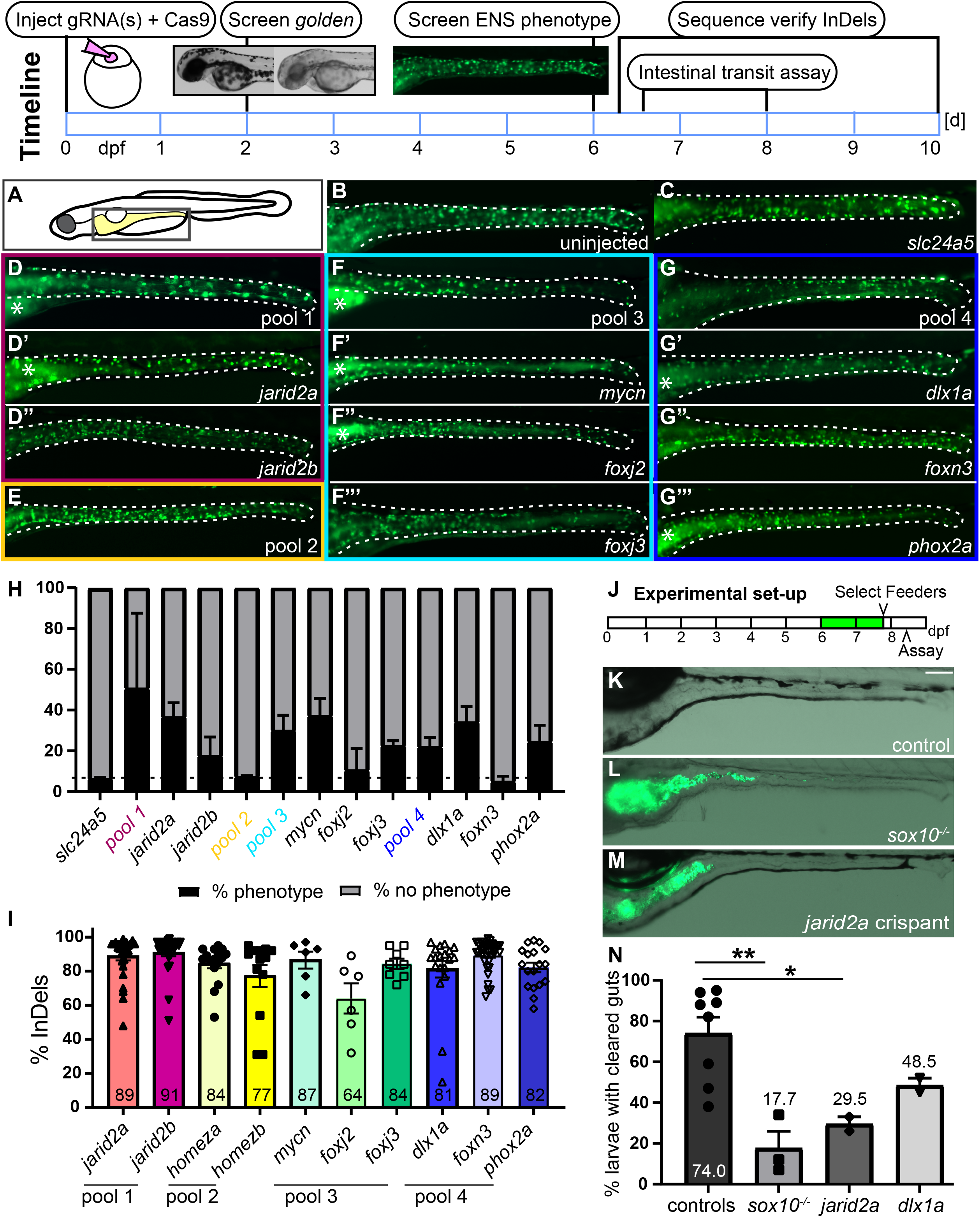
F_0_ CRISPR screen identifies new regulators of ENS neurogenesis and function. In 10 days, one set of gRNAs is phenotypically analyzed and Insertions and/or Deletions (InDels) at the target site are validated. This involves (1) injections of individual or pools of target guide RNA(s) + *slc24a5* gRNA + Cas9 protein, (2) screen for reduction of pigmentation (shown in the inset: wildtype left, *golden* phenotype right) at 2 dpf, (3) screen for ENS phenotypes at 6 dpf, (4) performing intestinal transit assay, and (5) sequence verify InDels at the target site. (**A**) Schematic of zebrafish larvae, the boxed area indicates area shown in B-G’’’. (**B**) Uninjected control and (**C**) F_0_ *slc24a5* crispants show wildtype levels of GFP^+^ ENS neurons (green) at 6 dpf. (**D, F, G**) Examples of 6-d-old F_0_ crispants with fewer GFP^+^ cells targeting *jarid2a* + *jarid2b* (pool 1), *mycn* + *foxj2* + *foxj3* (pool3), *dlx1a* + *foxn3* + *phox2a* (pool 4), but not (**E**) *homeza* + *homezb* (pool 2). Examples of F_0_ crispants targeting each gene individually from the three pools with ENS phenotypes show fewer GFP^+^ cells in *jarid2a* (**D’**), *mycn* (**F’**), *foxj2* (**F’’**), *foxj3* (**F’’’**), *dlx1a* (**G’**), and *phox2a* (**G’’’**), but not *jarid2b* (**D’’**) and *foxn3* (**G’’**) crispants. (**H**) Average percentage of crispants with and without reduced ENS phenotypes for each pool and each individual gene (≥ 2 experiments, % shown as mean +/− SEM). The dashed line shows the percent phenotypes for the *slc24a5* control crispants. (**I**) F_0_ crispants for each gene indicated show a high percentage of InDels in PCR amplicons of the target area (≥ 2 experiments, % shown as mean +/− SEM). (**J**) Experimental set-up for the intestinal transit assay. Larvae are fed fluorescently labeled food at 6 and 7dpf. On day 7, 6h post feeding, larvae that have eaten food are selected (“feeders”) and placed in a new dish without food. 16h later, clearance of labeled food from the gut is determined. In *sox10* mutant larvae (**L**), fluorescently labeled food (green) is still visible in guts compared to completely cleared guts of control larvae (**K**). In F_0_ crispants targeting *jarid2a* (**M**), fluorescently labeled food (green) remains visible. (**N**) Quantification of gut clearance (≥ 2 experiments, % shown as mean +/− SEM) in controls, *sox10* mutant larvae, *jarid2a* crispants, and *dlx1a* crispants. *sox10* mutant larvae and *jarid2a* crispants have significantly lower percentages of larvae with cleared guts. *dlx1a* crispants show a trend of fewer larvae with cleared guts compared to control larvae. B-G’’’. Whole-mount side views of 6 dpf zebrafish larvae in the area indicated in the schematic. K-M. Whole-mount side views of 8 dpf zebrafish larvae in the area indicated in the schematic. Asterisks: autofluorescence in the gut lumen; dashed line: gut outline; dpf: days post fertilization.

Next, we examined ENS function in F_0_ *jarid2a* and *dlx1a* crispants exhibiting reduced ENS neurons. An intestinal transit assay that measures if food travels through the gut ^8^ was utilized to assess gut function (Fig. 1J). In *sox10* mutant larvae and *jarid2a* crispants, the average clearance rate is significantly lower compared to controls (Figure 1K-N). In *dlx1a* crispants, we saw a trend of reduced average clearance rate compared to controls (Figure 1N).

## Discussion

This F_0_ CRISPR/Cas9 genome editing screen is a rapid, scalable, and inexpensive approach to functionally test candidate genes for a role in ENS neurogenesis and function. We find that (1) F_0_ crispants show changes in ENS neuron numbers and phenocopy known stable mutant phenotypes, (2) our screen identifies 5/10 new potential regulators of ENS neurogenesis, and (3) evaluating ENS function in a subset of these genes shows decreased intestinal transit in F_0_ crispants.

Of 10 candidate genes functionally tested in this CRISPR screen, five genes were determined to reduce ENS neuron number, suggesting that they are regulators of ENS neurogenesis. Our findings have revealed additional information on genes studied in mice for their role in regulating ENS neurogenesis. For example, in mouse *Phox2a* mutants, early ENS development was not affected and later ENS neurogenesis was not analyzed ^9^. In contrast mouse *Dlx1* mutants have normal ENS neuronal density but slowed intestinal gut motility ^10^.

In zebrafish, F_0_ CRISPR genome editing screens of candidate genes are effective approaches for prioritizing genes to generate stable mutant lines and subsequently analyze phenotypes in-depth ^4^. Determining the most promising targets among a large list of candidate genes in F_0_ crispants is crucial since the establishment of homozygous mutant phenotypes requires a considerable amount of resources and time (approximately 9 months). Notably, F_0_ screens in zebrafish are advantageous over other model systems due to the accessibility for manipulation and visualization in large numbers during early developmental stages. In our F_0_ screen, we analyze loss-of-function effects on ENS neuron numbers and intestinal function in live zebrafish larvae within 10 days allowing for functional evaluation of 16 gRNA pools at 3 genes per pool in 3 weeks. Our screening parameter of reduced ENS neuron numbers indicate a role in regulating ENS neurogenesis ^3, 4^. The intestinal transit assay is a fast and cost-effective measure of impacts on gut motility, a process connected to gut phenotypes in humans, such as constipation or diarrhea ^3, 4, 8^. The screening process is scalable due to the option of multiplexing candidate gene pools: we have injected gRNA pools to target up to 3 genes + *scl24a5* control per injection for more efficient screening. If few genes among candidates are expected to cause phenotypes, more genes can be combined in a pool with up to 8 genes ^7^. This screening approach is amenable to incorporate additional aspects of ENS development, ENS function, or gut development.

## Material and Methods

### Zebrafish husbandry

All experiments were carried out in accordance with animal welfare laws, guidelines, and policies and were approved by Michigan State University Institutional Animal Care and Use Committee. The *Tg(phox2bb:EGFP)^w37^* line was maintained, bred, and staged as described previously ^6^.

### gRNA design and synthesis

Gene-specific gRNA sequences and corresponding genotyping primer sets were designed using Chop Chop (http://chopchop.cbu.uib.no/). The gRNA oligo was annealed to the scaffold gRNA primer containing a T7 RNA polymerase binding site and the dsDNA template amplified by Phusion High-Fidelity DNA polymerase (Thermo Fisher Scientific F530S). The resulting dsDNA template was purified with a DNA clean and concentrate kit (Zymo Research, D4013), in vitro transcribed with the MEGAscript^TM^ T7 kit (Invitrogen, AM1334), and purified with a RNA clean and concentrate kit (Zymo Research, R1013).

### Protein loading

A 2X Cas9 protein buffer solution was made containing 3200 pg/nl Alt-R^®^ S.p. Cas9 Nuclease V3, 500 μg (IDT, 1081059), 600 mM KCl, and 8 mM HEPES pH 7.5. Protein loading was achieved by mixing up to 2 μl gRNA (200 ng/ul) with the 2X Cas9 buffer to achieve a final injection solution of 100 ng/ μl (25 ng of each individual gRNA and 1600 pg/nl Cas9). The injection needle was calibrated to deliver 1 nl of the injection solution into the yolk/cell interface of one cell stage zebrafish embryos.

### Preparation of fluorescent tracer

Fluorescent green tracer was prepared as previously described ^8^ by mixing 100 mg of powdered larval feed (Larval AP100 100 micron, Zeigler Bros), 150 μl of yellow-green fluorescent 2.0 uM polystyrene microspheres (FluoSpheres carboxylate modified microspheres (2% solid solution, Invitrogen) and 50 ul of deionized water. The ingredients were mixed together on a watch glass to form a paste and then spread into a thin layer. The mixture was allowed to dry at room temperature in the dark. Once dry, the mixture is scraped off and crushed into a fine powder between 2 sheets of weighing paper. The fluorescent powder was stored in an Eppendorf tube in the dark for up to 3 months.

### Intestinal Transit Assay

At 6 dpf embryos were sorted for reduced or normal ENS by fluorescence and transferred to a petri dish with embryo media in groups of 25 embryos. The intestinal transit assay was adapted from that previously described ^8^. Starting on 6 dpf, embryos were fed 2 mg of the fluorescent powder per dish. At 7 dpf embryos were transferred to fresh embryo media and fed 2 mg of fluorescent tracer. In the afternoon of 7 dpf embryos were screened for feeders by the presence of fluorescent food in the gut. The feeders were then transferred to a fresh petri dish with embryo media. On the morning of 8 dpf embryos were transferred to a fresh dish, anesthetized and examined for clearance of fluorescent food from the gut using the Zeiss Axio Zoom V16 stereo microscope.

### Image acquisition

Images were acquired using a Zeiss Axio Zoom V16 stereo microscope and ZEN software. Images were processed and analyzed using Photoshop 2020 (Version 21.0.2, Adobe Systems, Inc., San Jose, CA, USA) to adjust contrast and figures were assembled using Adobe Illustrator 2020 (Version 24.0.2, Adobe Systems, Inc., San Jose, CA, USA).

### Sequence analysis

Sequence analysis was performed on larvae after phenotypic analysis. PCR products containing the target site were amplified with gene-specific primers (Suppl Table 1). The percentage of Insertion and/or Deletions around the target site was determined using the free bioinformatics analysis tool from Synthego using the standard settings (https://www.synthego.com/).

### Statistical Analysis

To determine significant differences, we performed an unpaired t-test using GraphPad Prism 9.1.2.

## Acknowledgments

We want to thank Carrie Kozel, Theresa Gunn, and Taylor Lawrence for excellent fish care and Grant Kunzelman and Chuhao Nie for technical assistance. This work was supported by funds from Michigan State University (JG) and the 2019 AGA-Allergan Foundation Pilot Research Award in Irritable Bowel Syndrome (JG).

