## Supplementary material for "A rapid F0 CRISPR screen in zebrafish to identify regulators of neuronal development in the enteric nervous system": Suppl Fig 1

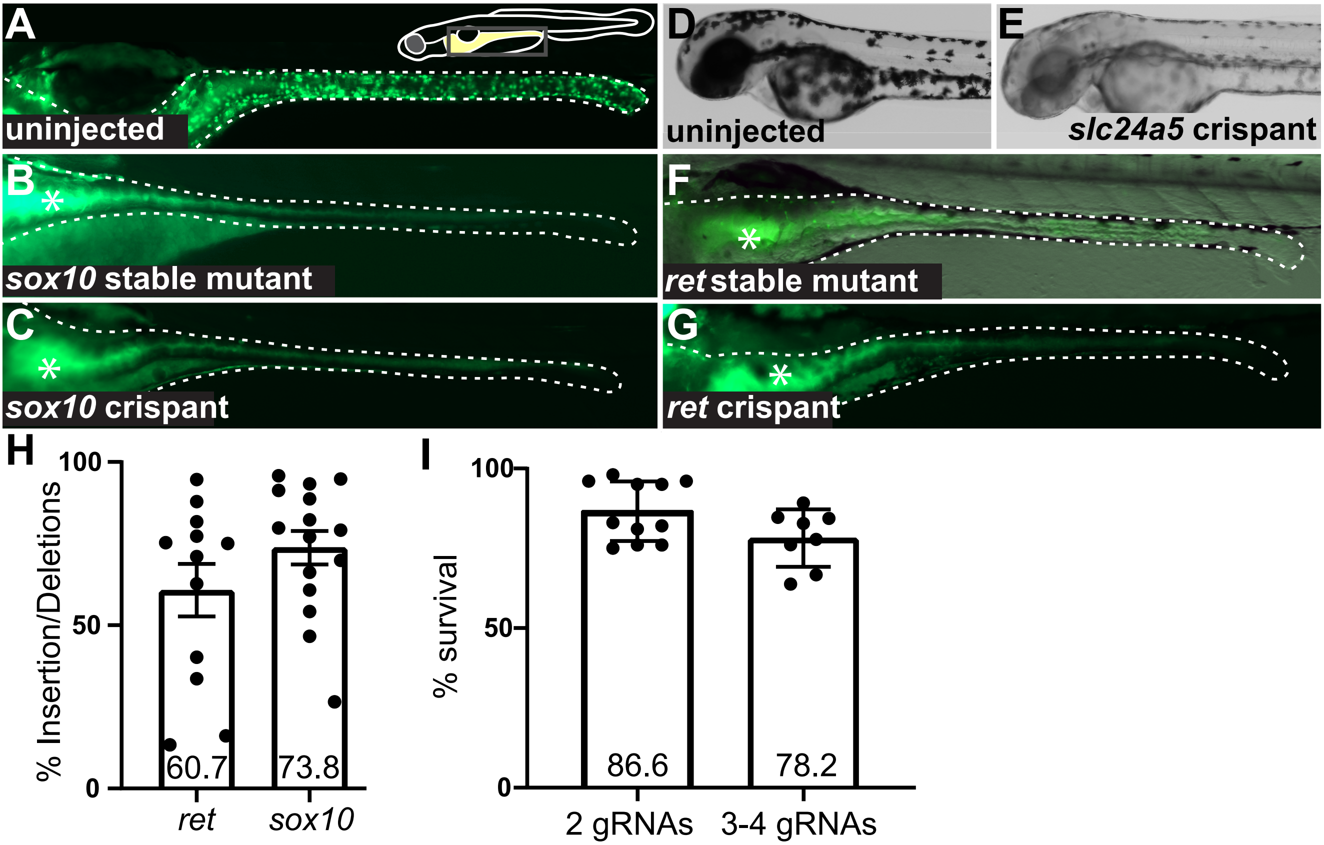
A
uninjected
B
sox10 stable mutant
C
sox10 crispant
D
uninjected
E
slc24a5 crispant
F
ret stable mutant
G
ret crispant
H
% Insertion/Deletions
100
50
0
60.7
73.8
ret
sox10
I
% survival
100
50
0
86.6
78.2
2 gRNAs
3-4 gRNAs

**Suppl. figure 1 Functional loss of *sox10* or *ret* in $F_0$ crispants fully recapitulates stable mutant phenotypes.** $F_0$ crispants phenocopy the lack of *phox2bb:GFP*$^+$ ENS neurons (green) seen in stable *sox10* or *ret* mutant larvae in comparison to wildtype-level ENS neurons in control larvae (**A**). Compare **B** and **C** (*sox10)* and **F** and **G** (*ret*). Functional loss of *slc24a5* (**E**) results in reduced pigmentation at 2 dpf compared to uninjected controls (**D**). (**H**) $F_0$ crispants have high percentages of Insertion and/or Deletions in PCR amplicons (≥ 1 experiment, % shown as mean +/- SEM). (**I**) Injected embryos show high survival at 2 dpf after injection of 2 or 3-4 guide RNAs (gRNAs, ≥ 8 experiments, % shown as mean +/- SEM). Whole-mount side views at 5 (A-C, F,G) or 2 (D,E) dpf. Asterisks: autofluorescence in gut epithelium; dashed line: gut outline; dpf days post fertilization.
