## Supplementary material for "A rapid F0 CRISPR screen in zebrafish to identify regulators of neuronal development in the enteric nervous system": Suppl Table 1

**Supplementary Table 1**

| | gRNA primer (5'-3') | Forward primer | Reverse Primer |
|---|---|---|---|
| control (*golden* phenotype) | | | |
| *slc24a5* _ex3_gRNA1 | aattaatacgactcactataGGTCTCTCGCAGGATGTTGCgttttagagctagaaatagc | ACTAAAACCACCATGTTTGTGT | AATTGTTTACCCAGAAATGCAG |
| **pool1** | | | |
| *jarid2a* _ex2_gRNA1 | attaatacgactcactata**g**TAAGTGAGAGGTAGAGCACCgttttagagctagaaatagc | TGTGTGTGTCTGCAGGATGATA | GTTGAGGGTCAAGGAAGTGAGT |
| *jarid2b* _ex2_gRNA1 | attaatacgactcactata**g**TGATGGGATGCCCTGGTCGGgttttagagctagaaatagc | CATTTGATGCTTTGCTGAGAAG | CTGTGCGCTCTTAAACTCCTTT |
| **pool2** | | | |
| *homeza* _ex2_gRNA1 | aattaatacgactcactataGAGACTCTGTAGCTGTAGGGgttttagagctagaaatagc | ACAGAACCAGCAAAGCTCTCTC | TCTCATCTTGTAAGCGCTGAAG |
| *homezb* _ex2_gRNA1 | aattaatacgactcactataGTTGTGGGTCGATGTGCCGGgttttagagctagaaatagc | CAGAGGCTAGGGAACAAGAAGA | GATGTGGGGTATGGAAATGAAT |
| **pool3** | | | |
| *mycn* _ex2_gRNA1 | aattaatacgactcactataGACTGGATAAGGGAAAACCAgttttagagctagaaatagc | CAAAGTCGTTCCTACTCCAACC | AGATTCTTACCAGAGTCGCTCG |
| *foxj2* _ex2_gRNA1 | aattaatacgactcactataGATGGCCATGGCGATCAGGGgttttagagctagaaatagc | TTCAGGGAAAGAAAGAGTGGAA | CGTATCGCTGATCCAAGTGTAA |
| *foxj3* _ex2_gRNA1 | aattaatacgactcactataGCCGCAGTTGACCATGCGGGgttttagagctagaaatagc | CTCCTTAGCTGCCACTCGAC | AGTGTCATTTTCTTCTTTGGCG |
| **pool4** | | | |
| *dlx1a* _ex2_gRNA1 | aattaatacgactcactataGAGGAGAGGTTCGTTTCAACgttttagagctagaaatagc | TTCTCCTTTTCTTTCTCCCTCC | GTAGCCTCTATTGCTTCCCTG |
| *foxn3* _ex2_gRNA1 | aattaatacgactcactataGCTTCGGTGACCCCGTGCTGgttttagagctagaaatagc | AGGAGATGGACCTGGCTCTT | GAGGTTTGCAGTTTGGGTTC |
| *phox2a* _ex1_gRNA1 | aattaatacgactcactata**g**CTCGTGCATGGCGGCGATGGgttttagagctagaaatagc | CTCAGAGCACCGGACACC | GAGCGGATCGGGTTGTACT |
| scaffold gRNA primer | GATCCGCACCGACTCGGTGCCACTTTTTCAAGTTGATAACGGACTAGCCTTATTTTAACTTGCTATTTCTAGCTCTAAAAC | | |
